# A comprehensive online database for exploring ~20,000 public Arabidopsis RNA-Seq libraries

**DOI:** 10.1101/844522

**Authors:** Hong Zhang, Fei Zhang, Li Feng, Jinbu Jia, Jixian Zhai

**Author notes:** Correspondence (J.Z.). These authors contributed equally to this work (Hong Zhang, Fei Zhang).

## Abstract

Application of Next Generating Sequencing (NGS) technology in transcriptome profiling has greatly improved our understanding of transcriptional regulation at genome-wide scale in the last decade, and tens of thousands of RNA-sequencing (RNA-seq) libraries have been produced by the research community. However, accessing such huge amount of RNA-seq data poses a big challenge for groups that lack dedicated bioinformatic personnel or expensive computational resources. Here, we introduce the Arabidopsis RNA-seq database (ARS), a free, web-accessible, and user-friendly to quickly explore expression level of any gene in 20,000+ publicly available Arabidopsis RNA-seq libraries.

---

In the last decade, RNA-sequencing (RNA-seq) has surpassed microarray to become the gold standard for gene expression profiling due to the continuous drop in sequencing cost and the development of easy-to-use library construction kits. To date, the *Arabidopsis* community has collectively released more than 20,000 RNA-seq libraries, with over 1,300 libraries deposited just in the first quarter of 2019 (Figure 1A). This vast resource is tremendously useful for all *Arabidopsis* researchers to study transcriptional regulation, tissue specificity, and developmental dynamics of genes they are interested in. However, accessing a large amount of RNA-seq data remains a big challenge for many groups that lack dedicated bioinformatic personnel or expensive computational resources.

**Figure 1.**
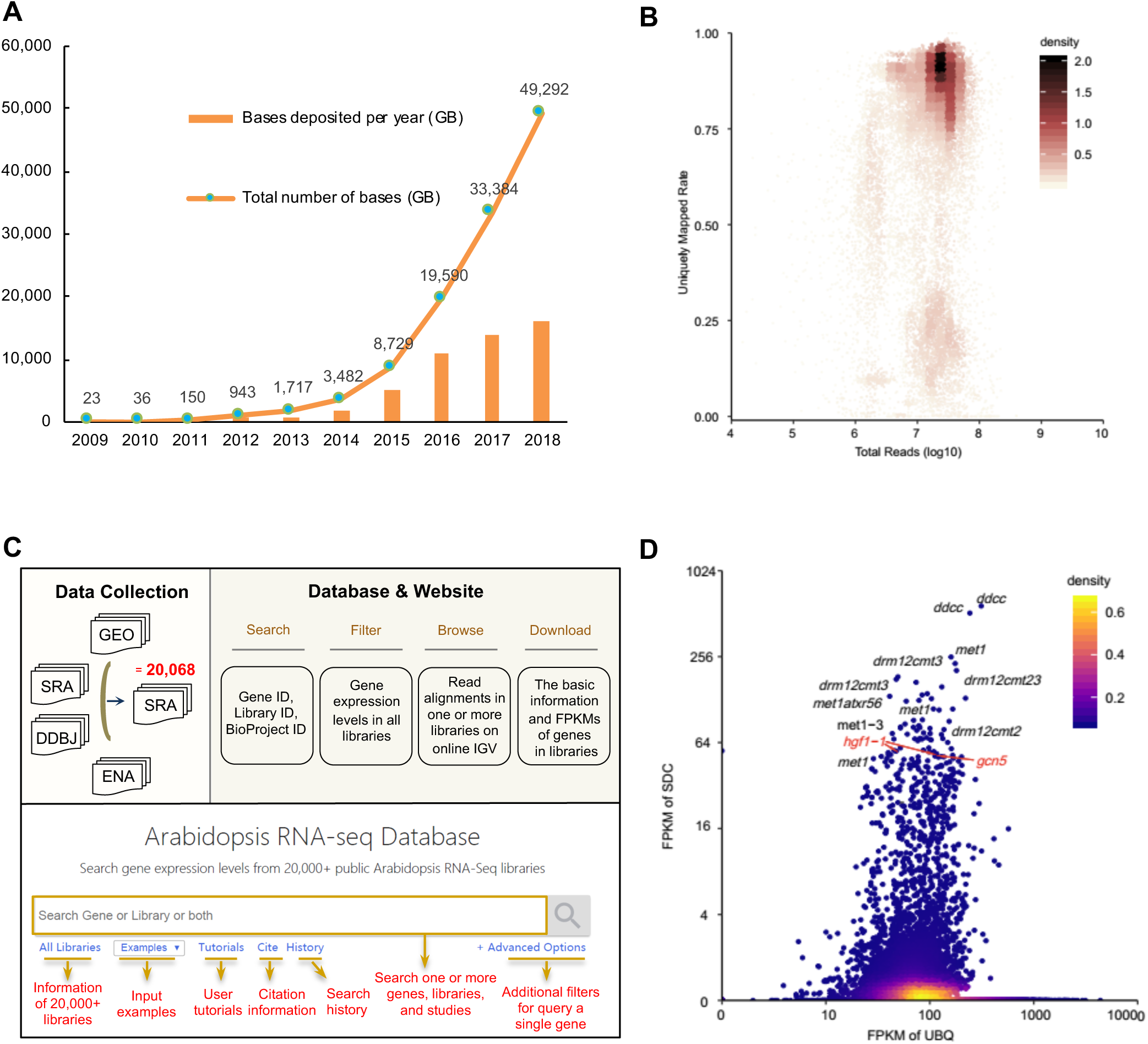
Overview of the Arabidopsis RNA-Seq Database. A. The number of sequenced bases per year from 2009 to 2018. X-axis represents the year of data generation, and y-axis is the number of sequenced bases in GB. B. Overall distribution of the percentage of uniquely mapped reads verses the total number of sequenced raw reads in 20,068 libraries. Each dot represents a library, and the color represents the density of the dots. C. Overflow for the construction of Arabidopsis RNA-Seq Database (ARS). A total of 20,068 publicly available Arabidopsis RNA-seq libraries were collected from GEO, SRA, DDBJ, and ENA databases, and processed with a unified pipeline. All genes and libraries-related information can be accessed via keyword-based searching on our ARS website (http://ipf.sustech.edu.cn/pub/athrna/). D. The expression levels of SDC in all libraries. A housekeeping gene UBQ is plotted on the x-axis. Some mutant libraries known roal in silencing of SDC is labeled black, and two potential novel players (*hgf1* and *gcn5* mutants) are labeled red.

Here, we present the Arabidopsis RNA-seq database (ARS, http://ipf.sustc.edu.cn/pub/athrna/) that integrates 20,068 publicly available Arabidopsis RNA-seq data deposited at Gene Expression Omnibus (GEO) (Barrett et al., 2013), Sequence Read Archive (SRA) (Kodama et al., 2012), European Nucleotide Archive (ENA) (Harrison et al., 2019) and DNA Data Bank of Japan (DDBJ) (Kodama et al., 2019) database (Table S1) before the end of March 2019. We downloaded raw data of all libraries and re-processed them with a standardized pipeline, mapped the reads to TAIR10 genome (Figure 1B), and calculated a normalized expression level in FPKM (Fragments Per Kilobase of transcript, per Million mapped reads) for all the 37,336 genes annotated in Araport11 (Cheng et al., 2017) in each library (Figure 1C).

ARS is a free, web-accessible, and user-friendly database which supports queries of gene IDs, library IDs, or BioProject IDs, or a combination of these to show specific genes in selected libraries. The search result is displayed in various forms, including data table, plots, and a built-in online IGV browser for convenient exploration. Taking the query results of AT2G17690 (*SUPPRESSOR OF DRM1 DRM2 CMT3, SDC*) as an example (Figure S1), the “Information” page shows the basic information of *SDC*, including the statistics of the maximum, minimum, median and mean value of FPKMs in all libraries, and the details of the SDC, such as locus type, alias, symbol, genome coordinate and gene direction (Figure S1A). The “Data Table” page presents the FPKM values of *SDC* in all ∼20,000 libraries, with library information such as the sample name, project, ecotype, genotype, tissue, release date of each library (Figure S1B). The “Data Plot” visualizes the FPKM values of *SDC* for easy comparison across multiple libraries (Figure S1D). ARS also integrates an online Integrative Genomics Viewer (IGV) (Robinson et al., 2017) to browse read coverage on *SDC* locus in selected libraries (Figure S1C).

Furthermore, our database can be used to quickly perform *in silico* screening among 20,000+ libraries to identify specific genotypes, conditions, or tissues that exhibit altered expression of any gene of interest. Still taking *SDC* as an example, which is a classic marker gene whose normal expression is strictly restricted to endosperm but can be found in somatic tissues when transcriptional gene silencing is defective (Henderson et al., 2008). A search of *SDC* in our database immediately identified RNA-seq libraries from mutants that are well-known components of the silencing pathway, such as *met1* and *drm12cmt23*, and also discovered two sets of libraries related to epigenetic regulation but have not been previously reported to be involved in silencing of the endogenous *SDC* (Figure 1D). The first set is from two biological replicates of *collin1* mutant (*hgf1-1* rep1 and rep2) (Figure 1D, Figure S2), which encodes a scaffold protein required for the formation of Cajal body (Kanno et al., 2016), which colocalizes with the AGO4 and NRPE1 in the nuclear processing center and require for the full function of DNA methylation (Li et al., 2006; Pontes et al., 2006; Li et al., 2008). Another set of libraries are three biological replicates from the mutant of *GCN5*, a histone acetyltransferase (Figure S2), and reader proteins for acetylated histone have been shown to regulate SDC expression (Zhang et al., 2016). Therefore, ARS provides quick, sensitive, and high-throughput screening methods to help identify novel players using the “bigdata” method. This approach will continue to grow in efficiency as the number of RNA-seq libraries deposited in public domains increases.

Many excellent web-based resources have been developed for hosting and analyzing mRNA-seq data, such as the MPSS database (Nakano et al., 2006), the UCSC Genome Browser (Kent et al., 2002), Anno-J Browser (Lister et al., 2008), EPIC-CoGe Browser (Nelson et al., 2018), ePlant (Waese et al., 2017) and CoNekt (Proost and Mutwil, 2018). Compared to these existing resources that are designed for either hosting a single-project or multiple-projects, or exploring the spatial and temporal dynamics of gene expression that requires smaller number of microarray/RNA-seq libraries (ePlant with 1,385 microarray samples, CoNekT with 913 RNA-seq samples), ARS can quickly extract the abundance information of any gene from 20,000+ Arabidopsis RNA-seq libraries using a simple “Google-like” search, and also provide easy access to view these data via a built-in online IGV browser. With a rapidly growing number of RNA-seq libraries, we plan to update ARS regularly in the future.

## RNA-seq data processing

We used the search term “*((Arabidopsis thaliana[Organism]) AND “transcriptomic”[Source]) AND “rna seq”[Strategy]*” to collect libraries from GEO and other public databases. For data processing, we aligned raw reads to TAIR10 reference genome using HISAT2 version 2.0.5 (Kim et al., 2015) with parameters “-max-intron-length=5000 -k 1 -dta --n-ceil -L,0,0.15”, and removed the duplicate reads with Samtools rmdup version 1.4.1 (Li et al., 2009). FPKMs were calculated by StringTie version 1.3.3b with the parameter “-e -r -G” (Pertea et al., 2015).

## Supporting information

Table S1. The detailed information of all RNA-seq libraries collected from public databases.

## Acknowledgements

We thank all the research groups that contributed RNA-seq data to the public domain, and we apologize for not being able to cite all the related papers in the main text due to limited space. References for all libraries that we used are listed in Table S1. The group of J.Z. is supported by National Natural Science Foundation of China (31871234), the Program for Guangdong Introducing Innovative and Entrepreneurial Teams (2016ZT06S172), and the Shenzhen Sci-Tech Fund (KYTDPT 20181011104005).

## Author Contributions

H.Z. processed the data, H.Z., F.Z., and L.F. built the database and website, J.J. and J.Z oversaw the study. H.Z., F.L., and J.Z. wrote the manuscript

## Supplemental Figures

**Figure S1.**
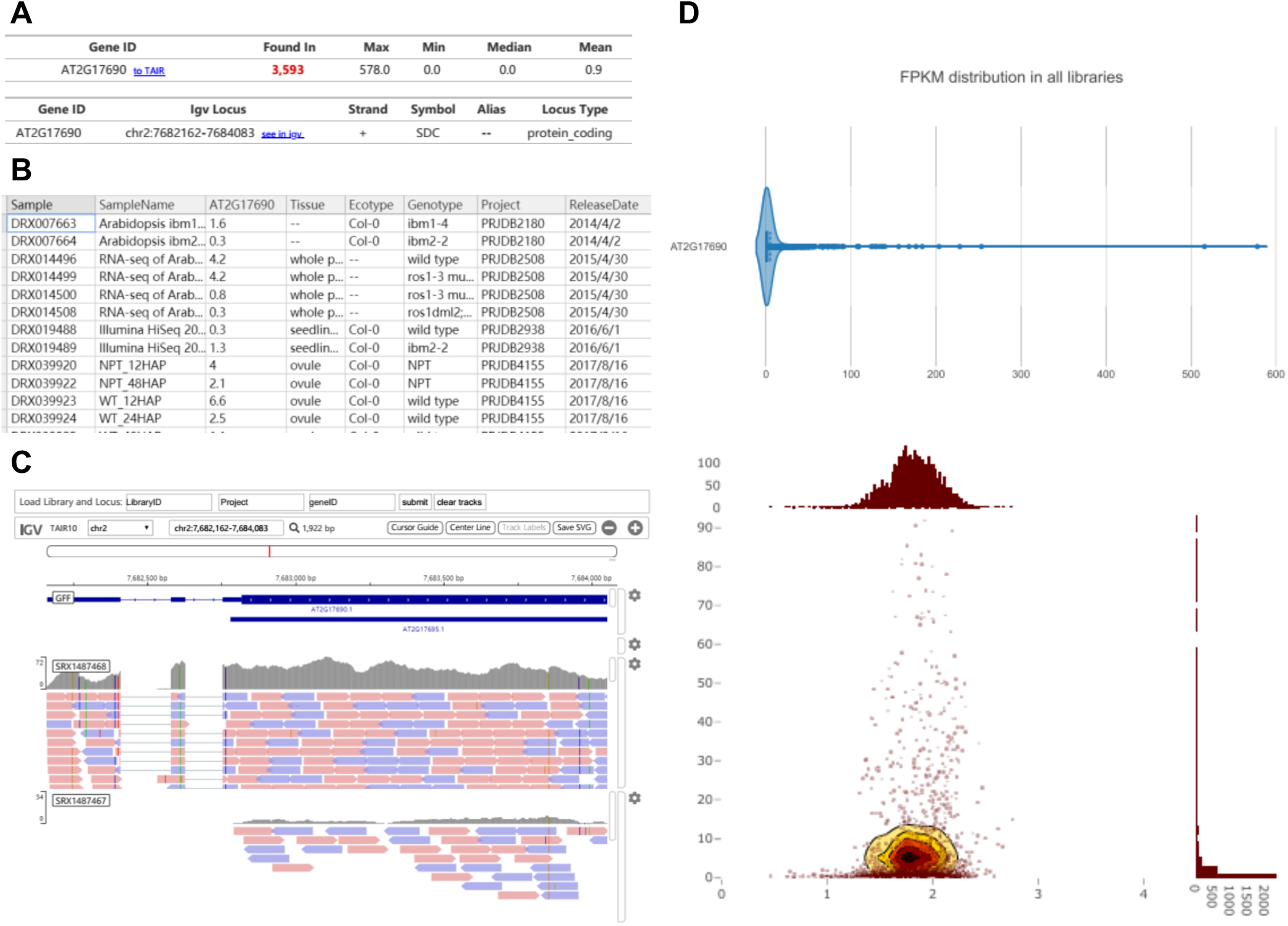
Example of a query using. SDC. A-D show the gene information, data table, data plot, and IGV browser, respectively.

**Figure S2.**
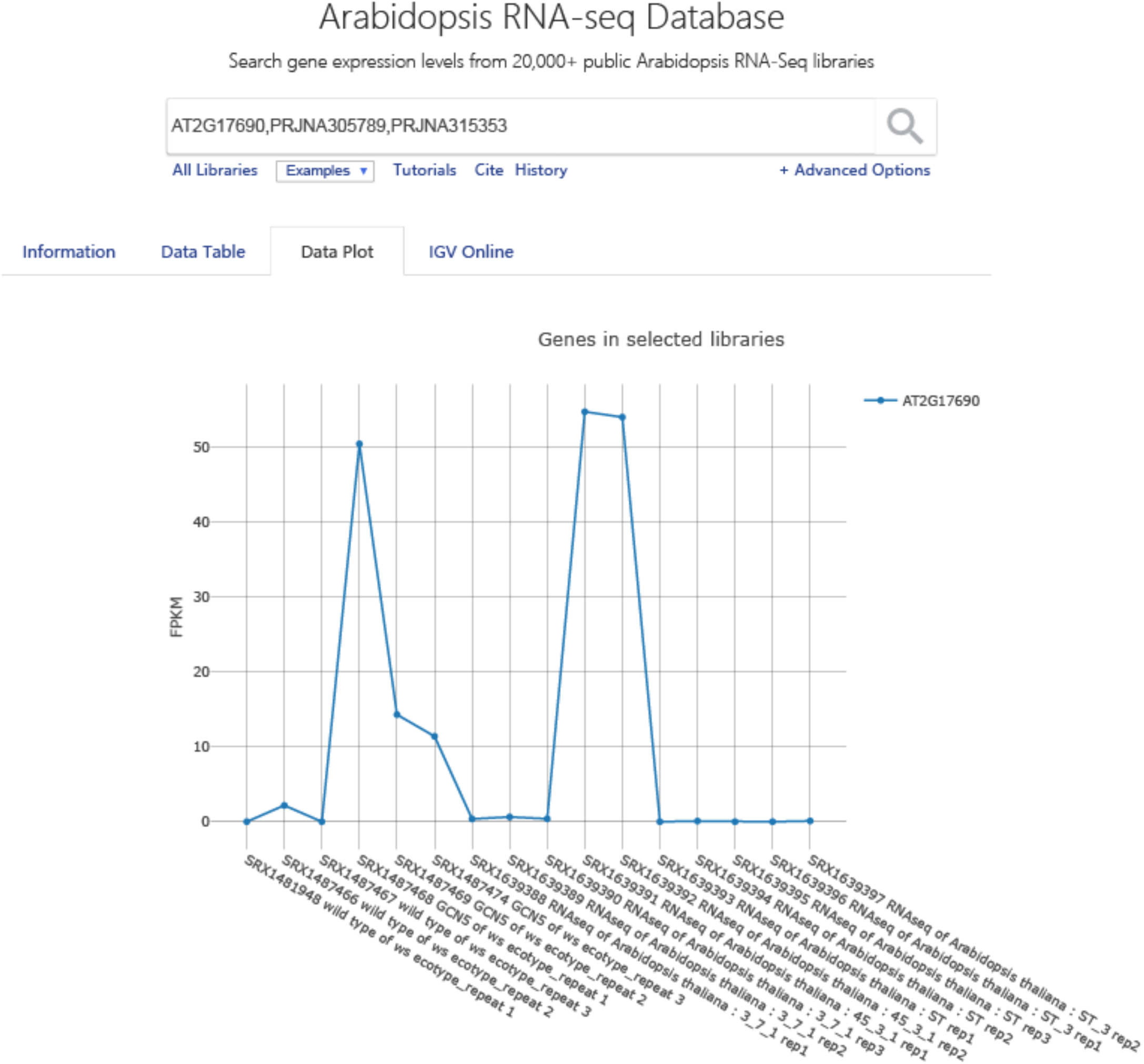
Expression level SDC in *hgf1* and *gcn5* mutants. The data plot page by searching AT2G17690 (SDC) and the two related BioProject numbers.

**Table S1. The detailed information of all RNA-seq libraries collected from public databases.**

